# LSD microdosing attenuates the impact of temporal priors in time perception

**DOI:** 10.1101/2023.04.14.536983

**Authors:** Renata Sadibolova, Clare Murray-Lawson, Neiloufar Family, Luke T. J. Williams, David P. Luke, Devin B. Terhune

## Abstract

Recent theoretical work embedded within the predictive processing framework has proposed that the neurocognitive and therapeutic effects of psychedelics are driven by the modulation of priors (Carhart-Harris & Friston, 2019). We conducted pre-registered re-analyses of previous research (Yanakieva et al., 2019) to examine whether microdoses of lysergic acid diethylamide (LSD) alleviate the temporal reproduction bias introduced by priors, as predicted by this theoretical framework. In a between-groups design, participants were randomly assigned to one of four groups receiving LSD (5, 10, or 20 μg) or placebo (0 μg) and completed a visual temporal reproduction task spanning subsecond to suprasecond intervals (0.8 to 4 sec). Using mixed-effects modelling, we evaluated the impact of the treatment group, and of the overall history of stimulus intervals (*global* priors) and the local stimulus history (*local* priors), weighted by their respective precision weights (inverse of variance), on temporal reproduction. Our principal finding was that the precision-weighted local priors and their precision weights reduced the under-reproduction bias observed under LSD in the original research. Furthermore, controlling for the precision-weighted local prior eliminated the reduced temporal reproduction bias under LSD, indicating that LSD microdosing mitigated the temporal under-reproduction by reducing the relative weighting of priors. These results suggest that LSD microdosing alters human time perception by decreasing the influence of local temporal priors.

## Introduction

Over the past decade, there has been a renewed interest in the neurocognitive effects of psychedelics, with recent clinical and basic research demonstrating this trend (Calder & Hasler, 2022; Carhart-Harris & Goodwin, 2017; Johnston et al., 2023; Lewis-Healey et al., 2022; Nichols, 2016). One influential model explaining the effects of psychedelics is the “<u>Re</u>laxed <u>B</u>eliefs <u>U</u>nder P<u>s</u>ychedelics” (REBUS) model, (Carhart-Harris & Friston, 2019), which proposes that these substances relax the weights on expectations based on prior experiences (priors). Our perception is influenced by our past sensory experiences, and according to the REBUS model, the relaxation of these priors under psychedelics can lead to higher thresholds for susceptibility to illusions and less perceptual bias (Carhart-Harris & Friston, 2019).

Here we investigated whether the bias in temporal reproduction may be moderated by LSD’s influence on temporal priors.

Information about the external environment processed via sensory channels often varies in quality and is combined with prior predictions for an optimal outcome to guide decision making (Friston, 2009; Körding & Wolpert, 2006; Raviv et al., 2012).

When priors and incoming information misalign, prediction errors arise, and the perceptual system strives to reduce the uncertainty. Precise priors are afforded higher weighting and override prediction errors but they are attenuated and updated when higher precision weights are afforded to incoming signals (Clark, 2013).

Psychedelics are thought to mitigate the influence of priors, thus yielding increased confidence in bottom-up information, with potential therapeutic benefits for psychiatric disorders hypothesized to be characterized by pathologically over- weighted priors (Cassidy et al., 2018; Powers et al., 2016; Teufel et al., 2015).

Recent research has supported this hypothesis, showing that psychedelics can reduce the prior precision weighting, leading to a more flexible and exploratory process of perception and cognition (Leptourgos et al., 2022; Muthukumaraswamy et al., 2013; Rajpal et al., 2022).

In addition to their effect on priors, psychedelics have been shown to modulate time perception (Coull, Morgan, et al., 2011; Kenna & Sedman, 1964; Wittmann et al., 2007; Yanakieva et al., 2019). For instance, Yanakieva et al. (2019) found that LSD reduced under-reproduction bias in their participants’ perception of stimulus durations compared to the placebo group (Fig. 2 in Yanakieva et al., 2019).

However, whether the under-reproduction could be attributed to the influence of temporal priors that was mitigated in the LSD group was not assessed. Accumulating research has shown that interval timing performance can be modelled as a form of Bayesian inference (Karaminis et al., 2016; Sadibolova & Terhune, 2022; Shi et al., 2013; Shi & Burr, 2016), thereby offering the possibility of scrutinizing the predictions of REBUS model through the analysis of the impact of psychedelics on temporal priors. In particular, Bayesian models of time perception posit that the brain forms temporal priors by extracting statistical patterns from the environment such as the mean duration of stimulus intervals (*global prior*, Acerbi et al., 2012; Jazayeri & Shadlen, 2010) and that temporal duration estimates gravitate towards the stimulus duration in previous trials (*local prior*; de Jong et al., 2021; Wiener et al., 2014).

These priors are combined with sensory evidence regarding the duration of a stimulus and can introduce temporal biases (Acerbi et al., 2012; Cicchini et al., 2012) that may be mitigated by administration of psychedelics.

In summary, by integrating the REBUS model into the predictive coding framework, it is possible to gain a more comprehensive understanding of the effects of psychedelics on time perception. Insofar as the pattern of findings by Yanakieva et al. (2019) conforms to the predictions of the REBUS model of the diminished impact of priors under psychedelics, we re-analyzed their data to investigate the possibility that their observations may be accounted for by global and local temporal priors interacting with the drug treatment. We predicted that the reduced impact of priors under psychedelics would remedy the temporal reproduction bias (i.e., the tendency for reproduced intervals to shift toward priors and away from objective interval durations).

## Materials and Methods

### Participants

This study involves re-analyses of previous data (Yanakieva et al., 2019); our analyses were preregistered on the Open Science Framework (https://osf.io/hfkjr).

The participants were 48 English-speaking adults, aged 55-75 (Mean age = 62.92, 21 female [44%], 27 male [56%]). Participants were randomly assigned to one of four groups (*n*=12) that received either LSD (5, 10, or 20 μg) or a placebo (0 μg). The removal of 2 multivariate outliers (one each from the 5 and 20 μg dose groups [see Yanakieva et al., 2019]) reduced the final sample size to 46.

### Experimental design

The data collection took place in an inpatient unit, adhering to standardized pre-screening and medical protocols (Yanakieva et al., 2019) as part of a larger pre- clinical trial on the safety and efficacy of microdose LSD (Family et al., 2020). The study used a randomized, double-blind, placebo-controlled design. The LSD solution was prepared in distilled water at a pharmacy on-site, and placebo groups were administered only distilled water, rendering the LSD and placebos indistinguishable to researchers and participants. The manufacturer of the drug product was Onyx Scientific Limited UK, to cGMP standards (Yanakieva et al., 2020).

Participants completed a temporal reproduction task once post-dose on the 4th day of dosing with the specific time of completion varying across participants (Yanakieva et al., 2019). Each trial of the task consisted of a fixed 750 ms cue “*memorize*”, a blank jittered interval (425–650 ms), and a target stimulus interval (blue circle [80 × 80 pixels; ∼2 cm in diameter], on a 1280 × 800 pixel-monitor) of varying duration (800, 1200, 1600, 2000, 2400, 2800, 3200, 3600, or 4000 ms). The stimulus was followed by a fixed 500 ms blank interval and a response cue “*reproduce*”, at which point participants responded by holding down the space bar to reproduce the stimulus interval. A blue circle co-appeared with this response and remained on the monitor until the spacebar was released. Trials were separated by a fixed 500 ms blank interstimulus interval. Participants completed one practice block, followed by four experimental blocks of 27 trials (108 trials total).

### Statistical analysis

We first removed bivariate outlier responses across stimulus intervals for each participant (*M*=8.37%, *SD*=2.95%, range: 1.85-14.81%) identified with the median absolute deviation (MAD) method implemented in the robust correlation toolbox (Pernet et al., 2013) in MATLAB (MathWorks, Natick, USA).

Data were analyzed with mixed effects modelling using the *lme4* package (Bates et al., 2015) in R (R Core Team, 2021). Our experimental design is characterized by a hierarchical structure (Figure S1 in supplemental materials) with *participants* (categorical; 46 levels) nested in the *dose* (continuous) and the dose nested in *cohorts* (categorical; 4 levels). As per Yanakieva et al. (2019), we created another dichotomous variable *drug* comprised of placebo and LSD levels which was included in our models instead of the *dose*. The models included additional continuous variables *stimulus interval* and *prior*. We further distinguished between different types of priors. The global prior was calculated at each trial (starting from trial 3) as the arithmetic mean of all preceding intervals. There were three local priors: the last preceding stimulus interval (*n-1 prior*), and the arithmetic means of the last preceding two intervals (*n-2 prior*) and three intervals (*n-3 prior*). The continuous variables (reproduced intervals, stimulus intervals, and priors) were mean-centered and the categorical variables (*drug* and *cohort*) were dummy coded.

Mixed-effects models were fitted using restricted maximum likelihood (REML) and the nonlinear “*nlopt_ln_neldermead”* optimizer (Wächter & Biegler, 2006) with tighter tolerance values (*1^e-12^*). Model fit improvement was determined by a change in the log-likelihood (increases with goodness of fit), and log-likelihood derived Bayesian Information Criterion (BIC; Schwarz, 1978) and Akaike Information criterion (AIC; Akaike, 1974) (both decrease with superior goodness of fit). We refrained from using the ꭕ^2^-distributed log-likelihood ratio test for model comparison as it’s been reported to produce anti-conservative *p*-values (Pinheiro & Bates, 2000). Instead, we applied Kenward and Roger’s approximation of degrees of freedom for *F*-test *p*- values and we computed the *p*-values for fixed effects parameters with Kenward and Roger’s method, as implemented in the “*pbkrtest”* and “*afex*” R packages (Halekoh & Højsgaard, 2014; Kuznetsova et al., 2017; Singmann et al., 2018). Furthermore, we employed the type III sums of squares method, which is suitable for unbalanced designs (Keppel & Wickens, 2004), due to sample size differences between the drug and placebo groups. To generate reliable estimates of uncertainty, we used the “*bootMer*” function to compute the bootstrapped 95% confidence intervals with 100 iterations. To further investigate the interaction effects, we conducted additional post hoc tests such as t-tests.

In order to determine the random-effects structure for the mixed-effects model, we began by generating null (’empty’) models (Quené & van den Bergh, 2004) comprising distinct random-effects structures (Barr et al., 2013), as outlined in Section S2 of the Supplementary Materials, which also includes the model diagnostics. The final random-effects structure of the model with the lowest BIC and AIC included correlated by-subject within-dose within-cohort intercepts and slopes for stimulus intervals. It was applied in all subsequent mixed-effects model analyses with fixed-effects parameters (stimulus intervals, drug, priors, and interactions thereof). Different priors were evaluated in separate models.

#### Unregistered prior precision-weights

In our registered analyses, we conceptualized priors as stimulus history, but did not take into account the crucial role of their precision weighting before their integration with sensory evidence (likelihood) (Petzschner et al., 2015; Sadibolova & Terhune, 2022; Shi et al., 2013; Shi & Burr, 2016). To rectify this omission, we replicated our registered analyses using prior precision weights, with higher values reflecting lower uncertainty for prior evidence, and precision-weighted priors. We calculated the precision weights as the inverse of variance for the prior normalized by the sum of inverse variances of the prior and likelihood (Petzschner et al., 2015). The initial prior precision at each trial was the inverse of the variance of all preceding trials (*global* priors), the preceding two trials (*n-2* priors), and the preceding three trials (*n-3* priors). We computed the likelihood variance for each trial by measuring the variability in reproduced intervals after regressing out the influence of priors, following the method introduced by Aston and colleagues (2021). For global priors, we regressed the preceding responses on the stimulus intervals and divided the variance of the residuals by the squared slope to remove the central tendency bias (Aston et al., 2021) whereas for the local priors, we regressed the reproduced intervals on each prior itself. As with the prior precision estimates, the initial likelihood precision was computed as the inverse of the likelihood variance.

## Results

### LSD attenuates performance bias in temporal reproduction

Our first expectation was for the mixed effects model to replicate the difference in reproduced supra-second intervals between the placebo and LSD treatment groups (Yanakieva et al., 2019). Our analyses replicated the reduced under-reproduction of stimulus intervals longer than 2 s (Yanakieva et al., 2019) in the LSD relative to the placebo condition (Figure **1**). In particular, the null model including the by-subject within-dose within-cohort intercepts and slopes for intervals (AIC=4020, BIC=4050) improved with the inclusion of Stimulus interval as a fixed-effects parameter (*Baseline model*), *F*(1,44.99) = 829.49, *p*<.001, *AIC*=3885, *BIC*=3922. The fit was further improved with the inclusion of Drug and Stimulus interval ⃰ Drug fixed-effects predictors (*Replication model*), *F*(2,42.66) = 4.91, *p*=.012, *AIC*=3880, *BIC=*3929 (Table **1**). The Stimulus interval, *β*=.64 (95% CI [.54, .73]), *SE*=.05, *t*=13.44, *p*<.001, Drug, *β*=.24 (95% CI [.08, .39]), *SE*=.10, *t*=2.37, *p*=.022, and Stimulus interval x Drug interaction, *β*=.17 (95% CI [.07, .27]), *SE*=.06, *t*=3.04, *p*=.004, predictors were all statistically significant. These results suggest a .64s increase in reproduced intervals for each 1s stimulus increment irrespective of the drug condition, reflecting a shallower slope than would be expected with veridical performance. However, relative to the placebo condition, the reproduced intervals under LSD were longer by .24s on average and they rose by .17s per 1s actual increment across stimulus intervals. Cumulatively, these results demonstrate that LSD induced a decrease in the temporal under-reproduction bias, resulting in a shift towards a more accurate and veridical performance.

**Figure 1.**
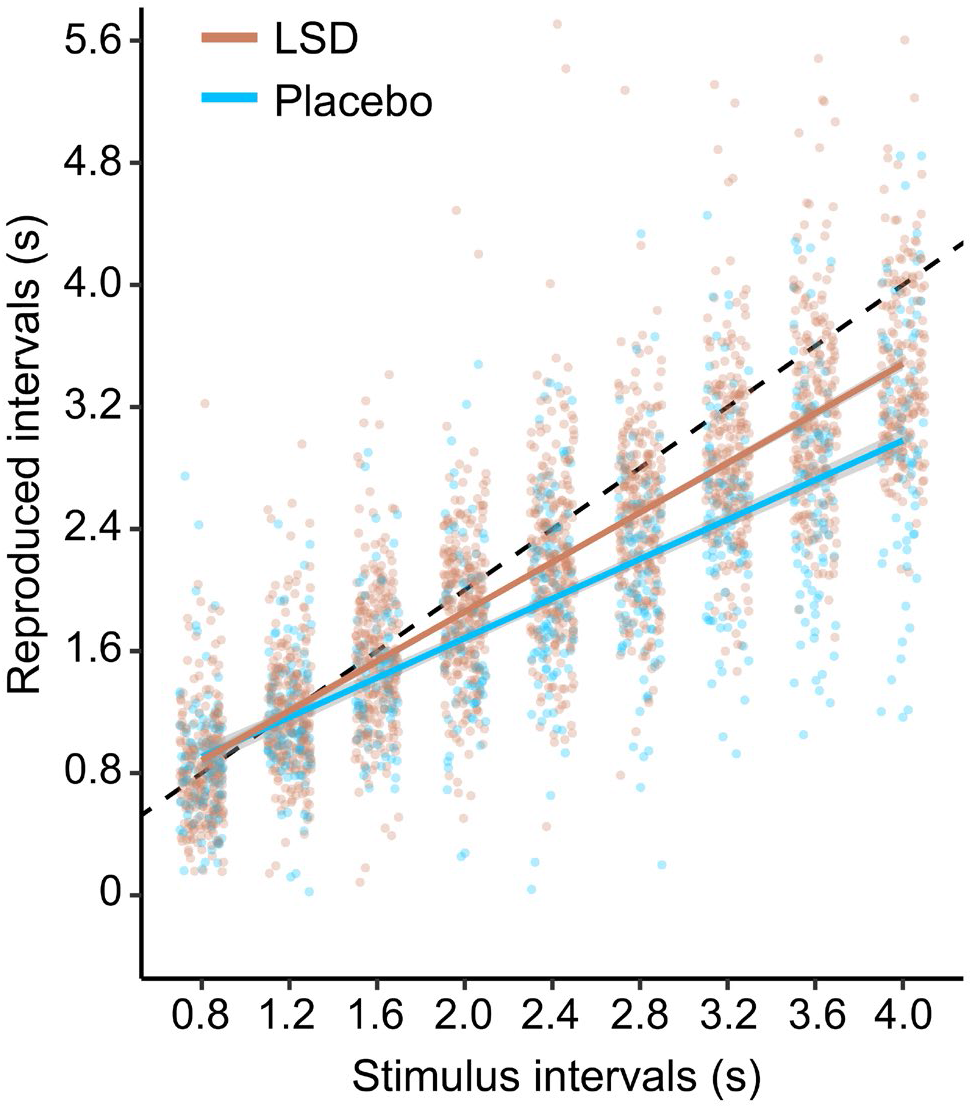
Temporal reproduction of stimulus intervals in LSD and placebo groups. The black dashed line represents veridical performance. Markers represent individual reproduced intervals. Shaded error bars are standard error smoothed with the linear *geom_smooth* function in R.

**Table 1.**
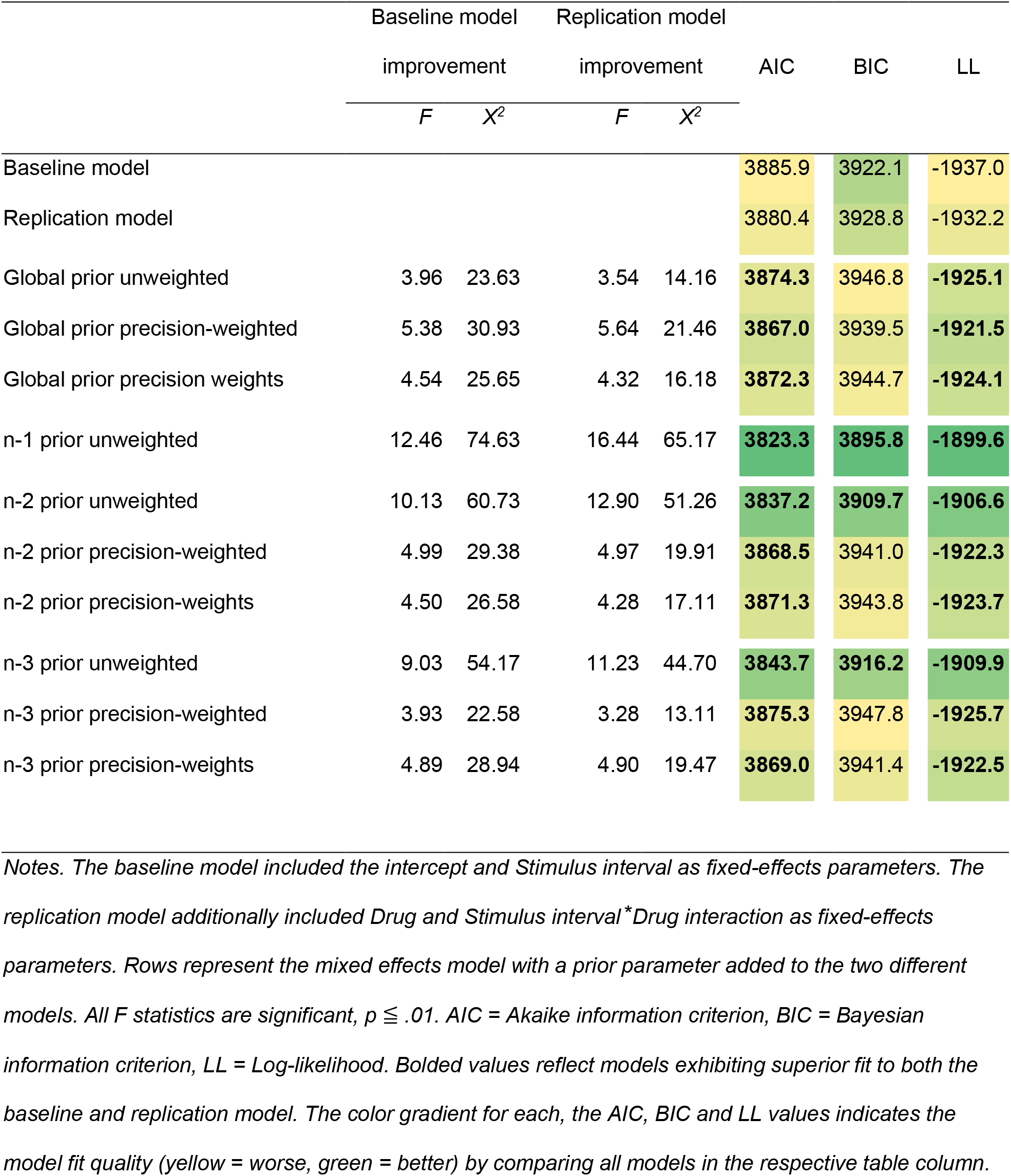
Comparison of baseline and replication mixed-effects models of temporal reproduction performance in placebo and LSD conditions.

|  | Baseline model |  | Replication model |  |  |  |  |
| --- | --- | --- | --- | --- | --- | --- | --- |
|  | improvement |  | improvement |  | AIC | BIC | LL |
|  | *F* | *X*<sup>2</sup> | *F* | *X*<sup>2</sup> |  |  |  |
| Baseline model |  |  |  |  | 3885.9 | 3922.1 | -1937.0 |
| Replication model |  |  |  |  | 3880.4 | 3928.8 | -1932.2 |
| Global prior unweighted | 3.96 | 23.63 | 3.54 | 14.16 | **3874.3** | 3946.8 | **-1925.1** |
| Global prior precision-weighted | 5.38 | 30.93 | 5.64 | 21.46 | **3867.0** | 3939.5 | **-1921.5** |
| Global prior precision weights | 4.54 | 25.65 | 4.32 | 16.18 | **3872.3** | 3944.7 | **-1924.1** |
| n-1 prior unweighted | 12.46 | 74.63 | 16.44 | 65.17 | **3823.3** | **3895.8** | **-1899.6** |
| n-2 prior unweighted | 10.13 | 60.73 | 12.90 | 51.26 | **3837.2** | **3909.7** | **-1906.6** |
| n-2 prior precision-weighted | 4.99 | 29.38 | 4.97 | 19.91 | **3868.5** | 3941.0 | **-1922.3** |
| n-2 prior precision-weights | 4.50 | 26.58 | 4.28 | 17.11 | **3871.3** | 3943.8 | **-1923.7** |
| n-3 prior unweighted | 9.03 | 54.17 | 11.23 | 44.70 | **3843.7** | **3916.2** | **-1909.9** |
| n-3 prior precision-weighted | 3.93 | 22.58 | 3.28 | 13.11 | **3875.3** | 3947.8 | **-1925.7** |
| n-3 prior precision-weights | 4.89 | 28.94 | 4.90 | 19.47 | **3869.0** | 3941.4 | **-1922.5** |

*Notes. The baseline model included the intercept and Stimulus interval as fixed-effects parameters. The replication model additionally included Drug and Stimulus interval\*Drug interaction as fixed-effects parameters. Rows represent the mixed effects model with a prior parameter added to the two different models. All *F* statistics are significant, $p \leq .01$ . AIC = Akaike information criterion, BIC = Bayesian information criterion, LL = Log-likelihood. Bolded values reflect models exhibiting superior fit to both the baseline and replication model. The color gradient for each, the AIC, BIC and LL values indicates the model fit quality (yellow = worse, green = better) by comparing all models in the respective table column.*

### The influence of priors on temporal reproduction

In the next subsections (also see Tables **1** and **S2**), we describe how the inclusion of global and local priors improved the model already including the stimulus and drug predictors. Motivated by the proposal that psychedelics modulate cognition and perception by attenuating prior precision weighting (Carhart-Harris & Friston, 2019), we predicted that temporal priors would predict reproduced intervals (*Baseline model* improvement) and that this effect would differ between LSD and placebo conditions (*Replication model* improvement).

#### Global priors (unweighted)

We first considered the role of global priors calculated as the mean of previous stimulus intervals at each trial. The global prior distribution is conventionally centered at the mean interval range, and research has demonstrated its association with central tendency bias in time perception, resulting in an overestimation of short intervals and underestimation of long intervals (Acerbi et al., 2012; Cicchini et al., 2012). The impact of including global priors in the *Baseline model* and *Replication model* is inconclusive (Table **1**), as the AIC and Kenward-Roger’s test p-values indicate improvement whereas the BIC values greater by more than 10 points suggest deterioration (Burnham & Anderson, 2004). Further results (see Table **S1**) did not support a differential role of global priors in temporal reproduction of long intervals under LSD (3-way interaction), *β*=.01 (95% CI [-.46, .39]), *SE*=.21, *t*=.03, *p*=.98. These observations seemingly contradict our prediction that the influence of global prior on central tendency bias is reduced in LSD condition. Indeed, rather than diminishing the influence of priors, LSD appears to have produced an increased reliance on them that generalized across stimulus intervals. In the placebo condition, the reproduced intervals follow the pattern of the global prior (albeit with overall under-reproduction), whereas in the LSD condition, the slope is steeper (Figure **2a**), Drug ⃰ Global priors, *β*=.50 (95% CI [.02, .94]), *SE*=.22, *t*=2.22, *p*=.030.

**Figure 2.**
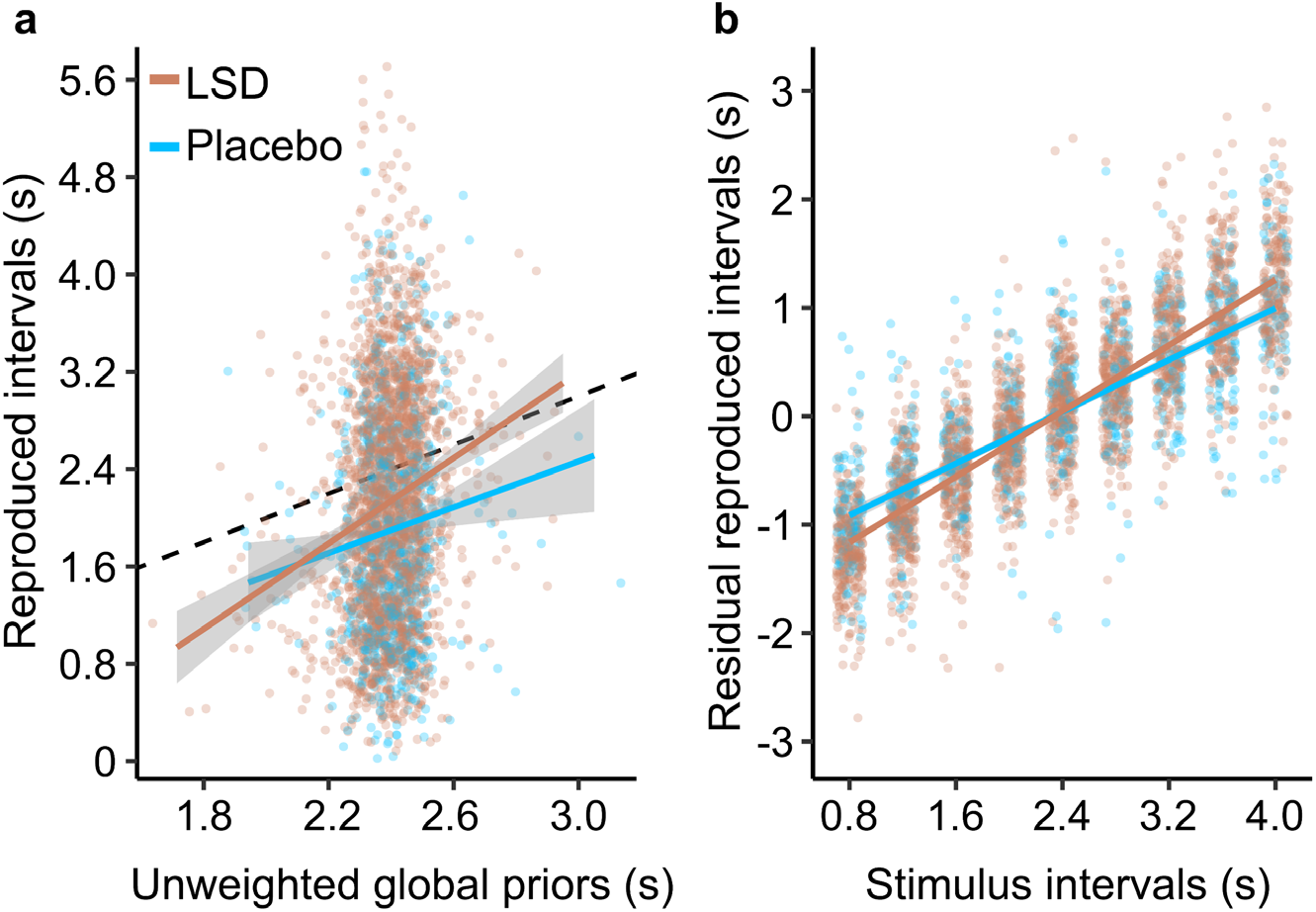
The effects of global priors on temporal reproduction performance in placebo and LSD conditions. **a.** Reproduced intervals (individual trial responses) as a function of unweighted global priors in each drug condition. The black dashed line represents the responses identical to global prior values. **b.** Residual reproduced intervals after removing the linear trend for global prior for each participant (see Figure 1 for raw reproduced intervals). Both panels: Markers are individual datapoints (trials). Shaded error bars are standard error smoothed with the linear *geom_smooth* function in R.

To test whether this effect contributed to reduced under-reproduction of longer stimulus intervals under LSD (Figure 1; Yanakieva et al., 2019), we fitted a model including Stimulus interval, Drug, and Stimulus interval ⃰ Drug fixed-effects to responses after removing trends for global priors for each participant (Figure **2b**). Although the resulting residual reproduced intervals continued to show a steeper increase with increasing stimulus intervals under LSD, β=.16 (95% CI [.063, .272]),

SE=.05, t=3.02, p=.004, the difference in under-reproduction bias across drug groups was no longer significant for three out of five previously significant long intervals (2.4, 2.8 and 4s, Holm-Bonferroni corrected p>.05). The mean reproduced intervals did not differ across group, β= -.01 (95% CI [-.039, .034]), SE=.02, t=-.25, p=.80. These analyses demonstrate that global priors partly shaped the pattern of the original observations (Yanakieva et al., 2019) although not via their reduced impact under psychedelics (Carhart-Harris & Friston, 2019).

#### Local priors (unweighted)

Our next analyses examined the roles of local priors in the impact of LSD on temporal reproduction. Incorporating local priors improved the model fit of both the *Baseline model* and *Replication model* (Table **1**). Reproduced intervals increased with longer preceding (*n-1*) stimuli, *β*=.07 (95% CI [.04, .10]), *SE*=.01, *t*=4.63, *p*<.001, and this effect varied across Stimulus intervals, Prior ⃰ Stimulus interval interaction: *β*=.03 (95% CI [-.01, .06]), *SE*=.01, *t*=1.99, *p*=.04. However, as with global priors, we did not find significant evidence for a Prior x Stimulus interval x Drug interaction, *β*= -.01 (95% CI [-.05, .02]), *SE*=.02, *t*=.52, *p*=.60, and we did not observe an interaction between the Prior x Drug group, *β*= -.01 (95% CI [-.05, .02]), *SE*=.02, *t*=.80, *p*=.42. Analyses using unweighted n-2 and n-3 priors exhibited similar patterns of results (see Supplementary Table **S1**). To summarize, our observations align with previous research (de Jong et al., 2021; Wiener et al., 2014) and suggest that reproduced intervals are reliably influenced by local priors not weighted by their precision, and that LSD does not moderate these effects.

#### Precision-weighted priors and precision weights

The foregoing approach can be expanded upon by considering the level of (un)certainty or precision of prior evidence, which determines how influential the priors are in shaping the responses (Petzschner et al., 2015; Sadibolova & Terhune, 2022). Therefore, we repeated the analyses but substituted priors with prior precision weights accounting for how impactful the priors are, and with priors weighted by their precision. We performed these analyses with all predictors *except* the n-1 priors, due to the inability to calculate their precision on trial-by-trial basis.

The impact of including precision weights and precision-weighted *global* priors in the *Baseline model* and *Replication model* remains inconclusive (Table **1**), as the AIC and Kenward-Roger’s test p-values continue to indicate improvement whereas the BIC values greater by more than 10 points suggest deterioration (Burnham & Anderson, 2004). Further results (Supplementary Table **S1**) indicate that the variation in reproduced time intervals between drug groups attributed to unweighted global priors (Figure **2**) lost statistical significance after accounting for global prior precision weighting. Altogether, these results do not furnish compelling substantiation for the global prior accounting for Yanakieva et al.’s (2019) findings.

Reproduced intervals decreased with increasing n-2 prior weights, *β*=-.15 (95% CI [-.26, -.05]), *SE*=.07, *t*=-2.19, *p*=.03, and n-3 prior weights, *β*=-.36 (95% CI [-.55, -.17]), *SE*=.09, *t*=-3.89, *p*<.01, indicating a greater local history bias for higher precision weights, as would be expected. The reduction in reproduced intervals driven by the prior precision weights occurred at a slower pace in the LSD condition compared to the placebo condition*, β*=.19 (95% CI [.07, .33]), *SE*=.08, *t*=2.38, *p*=.02 (n-2 precision weights * Drug), and *β*=.34 (95% CI [.11, .55]), *SE*=.11, *t*=3.07, *p*<.01 (n-3 precision weights * Drug). For n-2 precision weights, this effect was more pronounced for long stimulus intervals (n-2 precision weights * Drug * Stimulus), *β*=.19 (95% CI [.03, .35]), *SE*=.08, *t*=2.50, *p*=.01. Accordingly, the decrease in reproduced intervals (under-reproduction bias) as a function of n-2 priors adjusted by these precision weights was reduced for longer stimulus intervals in the LSD relative to the placebo condition, *β*=.07 (95% CI [.01, .13]), *SE*=.03, *t*=2.30, *p*=.02 (Table **S1**). In order to determine whether this effect accounts for the observation of reduced under-reproduction of long stimulus intervals in the LSD condition (Fig 2; Yanakieva et al., 2019), we fitted the model with Stimulus interval, Drug, and Stimulus interval *

Drug fixed-effects to residual reproduced intervals after removing the linear trend for precision-weighted n-2 priors for each participant. Our results suggest that the LSD-mediated reduced bias in the original study by Yanakieva and colleagues (2019) was indeed no longer significant after accounting for the influence of the precision-weighted n-2 priors, *β*=-.01 (95% CI [-.05, .03]), *SE*=.06, *t*=-0.29, *p*=.77, and the across-condition differences for individual long stimulus intervals were not statistically significant (Holm-Bonferroni corrected *p*s>.05). Taken together, our results suggest that reduced precision-weighting of local temporal priors underlies altered time perception in LSD (Yanakieva et al., 2019).

## Discussion

In this re-analysis of existing data (Yanakieva et al., 2019), we examined whether LSD alters time perception by modulating the impact of temporal priors on temporal reproduction. We predicted that the REBUS model’s theorized reduced impact of priors under psychedelics (Carhart-Harris & Friston, 2019) would remedy the temporal reproduction bias (i.e. the tendency for reproduced intervals to shift toward priors and away from objective interval durations). We found that the impact of *global* priors (unweighted by their precision) on temporal reproduction was more pronounced under LSD, contrary to the REBUS model predictions. However, the difference in reproduced intervals across drug groups was not fully eliminated when the influence of the global priors was controlled for, suggesting that they only partly explained group variation in Yanakieva et al.’s data. Moreover, the impact of the global priors on temporal reproduction was similar in both drug groups once they were weighted by their precision. By comparison, *local* prior precision and precision-weighted priors were associated with tempered under-reproduction of longer stimulus durations in the LSD group. Reproduced intervals decreased with increasing local prior precision weights, indicating a greater local history bias for higher prior precision, and the reduction in reproduced intervals driven by precision weights occurred at a slower pace in the LSD group compared to the placebo group. Further analyses showed that these effects accounted for the original observation (Yanakieva et al., 2019), given that the reproduced long intervals did not significantly differ across drug groups once the impact of the precision-weighted local prior was controlled for. These results suggest that altered temporal reproduction under LSD (Yanakieva et al., 2019) may be explained by local temporal priors in line with the proposal that psychedelics reduce the impact of priors (Carhart-Harris & Friston, 2019; Safron et al., 2020).

The REBUS model (Carhart-Harris & Friston, 2019) supports these findings by suggesting that psychedelics decrease the confidence of priors and reduce their constraining effect on the processing of incoming information (prediction errors). According to this model, the relaxation of priors is most evident at high levels of the processing hierarchy, such as those associated with the default-mode network, which are linked to self-hood, identity, and ego (Carhart-Harris et al., 2012).

However, the model also proposes that a wide range of functional levels will be impacted, including priors at intermediate levels of the processing hierarchy, albeit potentially with less conspicuous psychological effects (Carhart-Harris & Friston, 2019). Accordingly, our findings suggest that temporal reproduction under LSD is less influenced by temporal priors and therefore exhibits less bias under LSD. Our observations broadly align with previous research on the LSD-mediated reduced influence of priors on perception. For instance, LSD was found to reduce the brain’s ability to detect and respond to unexpected or deviant auditory stimuli, as indicated by the diminished amplitude of the mismatch negativity component in response to auditory stimuli (Timmermann et al., 2018). This was attributed to LSD reducing the precision of the brain’s internal models of expected input. Additionally, the reduced Kanizsa illusion, which relies on top-down predictions from higher visual areas, provides further evidence of the impact of psychedelics on priors (Kometer et al., 2011). Altogether, these findings provide valuable insights into the effects of psychedelics on sensory processing and prior predictions in shaping perception.

It has been repeatedly demonstrated that LSD and germane psychedelics induce pronounced alterations in perception of time, yet the neurocognitive and neurochemical mechanisms underlying these effects have not been fully understood (Altman, 1977; Aronson, 1959; Kenna & Sedman, 1964; Passie et al., 2008; Wittmann et al., 2007). Recent predictive processing theories, such as the REBUS model, offer a new perspective, proposing that psychedelics exert their influence on cognition by increasing the excitability of deep-layer pyramidal neurons that express 5-HT receptors (Carhart-Harris & Friston, 2019; Nichols, 2016; Safron et al., 2020). According to the model, the overly excitable neurons fail to synchronize, thereby reducing the influence of top-down priors and increasing the likelihood of the system being updated by unsuppressed ascending prediction errors. In this way, the REBUS model suggests that psychedelics afford a greater latitude for belief updating throughout the processing hierarchy (Carhart-Harris & Friston, 2019; Nichols, 2016; Safron et al., 2020) although the evidence for this neurocognitive mechanism and its neurochemical instantiation in temporal perception is currently underspecified.

Future research will benefit from investigating the neural mechanisms underpinning our observations to shed further light into how they align with these theoretical accounts.

Notably, the effects of LSD have been linked not only to its psychedelic properties via the activation of serotonin 5-HT receptors, but also to its psychostimulant properties via dopamine D1 and D2 receptors and dissociative properties via NMDA glutamate transmission (Marona-Lewicka & Nichols, 2007; Nichols, 2004; Passie et al., 2008). The dopamine system, in particular, has been widely implicated in altered temporal perception (for reviews, see Agostino & Cheng, 2016; Coull, Cheng, et al., 2011; Marinho et al., 2018). Interestingly, whereas the serotonin 5-HT agonism has been discussed in the context of psychedelic relaxation (down-weighting) of priors for its therapeutic effects in psychopharmacological models of psychotic hallucinations (Corlett et al., 2019; Haarsma et al., 2021; Rajpal et al., 2022), there is evidence for elevated striatal dopamine being associated with an over-reliance on priors (Cassidy et al., 2018). Furthermore, the LSD has been shown to decrease dopamine firing activity through 5-HT, D2 and TAAR1 receptors (De Gregorio et al., 2016; Marona-Lewicka & Nichols, 2007). Accordingly, an alternative interpretation of the present results is that an LSD-mediated reduction in striatal dopamine levels yielded the observed attenuation of precision-weighting of local temporal priors underlying the observed altered timing performance (Yanakieva et al., 2019). Further work is required to discriminate between these competing neurochemical interpretations. Additionally, the relationship between dopamine and serotonin systems in the effects of LSD on temporal perception warrants further investigation, as both systems may have opposing effects on temporal priors (Carhart-Harris & Friston, 2019; Cassidy et al., 2018).

Alternatively, our findings could be interpreted as resulting from increased information flow due to reduced thalamic gating under the cortico-striatal thalamocortical (CSTC) model (Preller et al., 2019). This model proposes that LSD, acting as a serotonergic agonist, diminishes the striatal influence on the thalamus, thus opening the thalamic filter. This interpretation is consistent with predictive processing accounts, which frame thalamic gating and increased “bottom-up” information flow to cortical areas in terms of the heightened precision of ascending prediction errors (likelihood) (Clark, 2013, 2016). This process reflects the gain on incoming information at the expense of priors, which in the case of temporal priors and their associated biases, leads to a reduction in reproduction biases under LSD, as observed in this study. Recent accounts of the CSTC model have further expanded its scope by including the claustrum, which is densely populated by 5-HT receptors and has been implicated in the temporal integration of cortical and thalamic oscillations for interval timing and working memory (Doss et al., 2022; Yin et al., 2016). Thus, future research would additionally benefit from investigating the roles of the thalamus and claustrum in mediating LSD’s effects on temporal priors in temporal perception.

Moving from the discussion of neurocognitive and neurochemical bases of the observed effects, it is worth exploring the significance of LSD micro-dosing. A typical full dose of LSD ranges from 75-150 µg and produces a diverse array of phenomenological effects such as hallucinations, ego dissolution and altered perception of space and time (Passie et al., 2008). By comparison, the phenomenological effects of LSD micro-dosing are minimal (Yanakieva et al., 2019). Therefore, unlike explicit psychoactive effects of LSD (e.g., mystical experiences) and putative therapeutic effects (e.g., antidepressant effects), which are plausibly shaped in part by participants’ expectancies (Burke & Blumberger, 2021; Butler et al., 2022; Olson et al., 2020), it is unlikely that participants expect to exhibit reduced temporal under-reproduction under LSD. This strongly suggests that the observed results are not attributable to explicit expectations, as has been suggested for other effects of microdose psychedelics (Kaertner et al., 2021). Future research could expand upon the present work by more stringently manipulating temporal priors under different LSD doses and investigating the role of specific neurochemical systems.

To summarize, our study has shed light on the intricate ways in which LSD impacts temporal reproduction by modulating local and global temporal priors. Specifically, we have demonstrated that even small doses of LSD can interact with local temporal priors, resulting in alterations in temporal reproduction. Our findings are underpinned by existing theoretical models based on the predictive coding framework, which offer a potential explanation for the observed effects. The results of this study pave the way for future research to further explore these underlying mechanisms. Overall, our findings align with the REBUS model of the effects of psychedelics on cognition (Carhart-Harris & Friston, 2019), suggesting that low doses of psychedelics have the potential to alleviate temporal biases imposed by temporal priors. These exciting results represent an important step in advancing our understanding of the effects of psychedelics on temporal perception.

## Acknowledgements

This work was supported by the Biotechnology and Biological Sciences Research Council Grant BB/R01583X/1 to DBT.

