## Supplemental Materials for "LSD microdosing attenuates the impact of temporal priors in time perception"

**S1. Hierarchical variable structure**

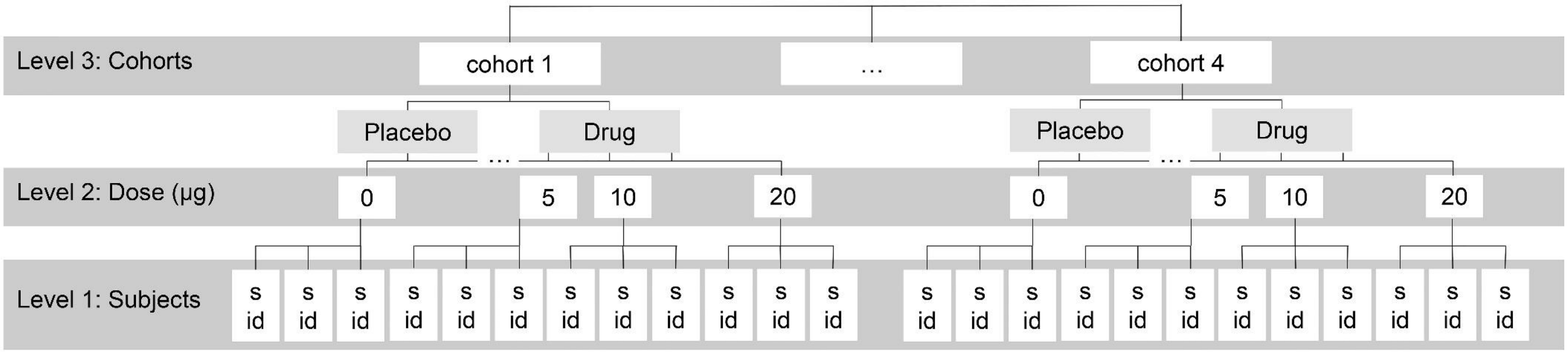


*Figure S1*. Multilevel structure of the experiment. The nested variables include participants and dose (additionally binarized as placebo vs. LSD). Subjects were additionally crossed with stimulus interval and priors (not included here).

### S2. Null model

We initially constructed a null ('empty') model (Quené & van den Bergh, 2004) including the maximal random-effects structure (Barr et al., 2013): (1) By-participant (nested in the dose and cohort) intercepts and slopes for the stimulus and their correlation, (2) By-dose (nested in the cohort) intercepts and slopes for the stimulus and their correlation, and (3) By-cohort intercepts and slopes for stimulus and their correlation. The resulting random-effects structure was further explored by comparing the intercept-only model, the slope-only model, and the correlated and uncorrelated random-effects models by means of fit statistics.

By removing the by-dose within-cohort and the by-cohort random effects from the model, we addressed the singular fits and arrived at a parsimonious mode.l (Matuschek et al., 2017). The by-subject within-dose within-cohort intercepts and slopes for stimulus intervals were retained. The by-subject within-dose within-cohort intercepts (Variance=.71, *SD*=.85) and slopes for stimulus (Variance=.61, SD=.78) were highly correlated (*r*=.96). However, the removal of either the random intercepts (*AIC*=5125.60, *BIC*=5143.70) or slopes (*AIC*=8290.20, *BIC*=8308.30) from the model would have substantially reduced fit (*AIC*=4020.40, *BIC*=4050.60) and was therefore avoided. The model's residual variance was .19 (*SD*=.43) and the RMSE = .43.

The null model assumptions were assessed with *check_model* function from performance R package (Lüdecke et al., 2021). The model fitted data well (Figure **S2a**), and the assumptions of normality of the random effects (Figure **S2b**), linearity and homoscedasticity of residuals (Figure **S2c**) were met. Although two participants showed larger residual variability (Figure **2e**) and a slight deviation from normality (Figure **2d**), they were not excluded from the analyses since almost all their datapoints fell below Cook's value of 1 (Figure **2f**).

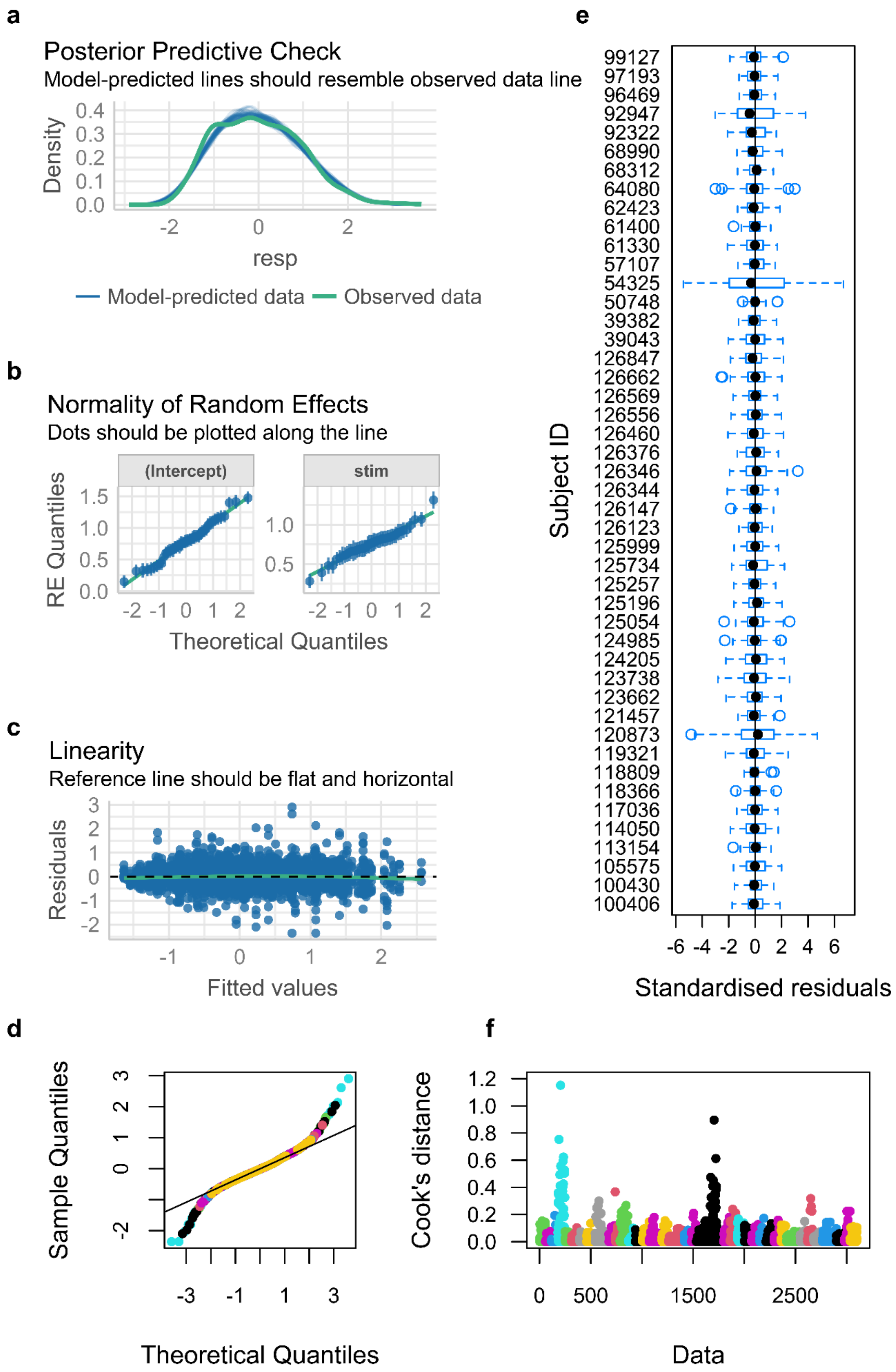


*Figure S2*. Null model diagnostic plots. **a.** Posterior predictive checks show similar density functions fitted to the observed and predicted reproduced intervals

(Lüdecke et al., 2021). **b.** Residual by quantiles plot showing normality of random effects structure. **c.** Standardized residuals plotted against the fitted values corroborate the assumptions of linearity and homoscedasticity. **d.** Normal QQ plot of model residuals, colour-coded by participant. **f.** Cook's distances for all data points colour-coded by participant. **e.** Boxplots of residual differences across participants.

#### S3. Linear mixed effects models including the priors

Supplementary table S2 provides a detailed overview of statistical results for all linear mixed effects models fitted in this study. These models share the random-effects structure (cf. Supplementary section S2), and the stimulus and drug and their interaction as the fixed-effects parameters. They each assess one of four priors (global prior and three local priors) and its interactions with the other two fixed-effects parameters. The models were constructed with the raw unweighted (U) and precision-weighted (W) priors, as well as the prior precision weights (wei) alone.

Table S1

*Linear mixed effects models fitted to reproduced intervals.*

**G(U)***stim*drug + (stim|cohort:dose:id)

| ***Fixed effects*** | ***b (SE)*** | ***t*** | ***p*** | |
|---|---|---|---|---|
| stim | 0.64 (0.05) | 13.45 | <0.01 | * |
| drug | 0.24 (0.10) | 2.37 | 0.02 | * |
| G(U) | -0.10 (0.20) | -0.52 | 0.61 | |
| stim*drug | 0.16 (0.06) | 2.90 | 0.01 | * |
| G(U) *stim | 0.07 (0.19) | 0.40 | 0.69 | |
| G(U) *drug | 0.50 (0.22) | 2.22 | 0.03 | * |
| 3-way inter | 0.01 (0.21) | 0.03 | 0.98 | |

| ***Random effects*** | | ***Variance (SD)*** |
|---|---|---|
| cohort:dose:id | intercept | 0.09 (0.29) |
| | slope (stim) | 0.02 (0.16) |
| | correlation | 0.58 |

**G(W)***stim*drug + (stim|cohort:dose:id)

| ***Fixed effects*** | ***b (SE)*** | ***t*** | ***p*** | |
|---|---|---|---|---|
| stim | 0.64 (0.05) | 13.84 | <0.01 | * |
| drug | 0.28 (0.10) | 2.74 | 0.01 | * |
| G(W) | 0.17 (0.10) | 1.75 | 0.08 | o |
| stim*drug | 0.15 (0.05) | 2.81 | 0.01 | * |
| G(W) *stim | -0.05 (0.06) | -0.79 | 0.43 | |
| G(W) *drug | 0.08 (0.13) | 0.65 | 0.52 | |
| 3-way inter | -0.06 (0.09) | -0.72 | 0.47 | |

| ***Random effects*** | | ***Variance (SD)*** |
|---|---|---|
| cohort:dose:id | intercept | 0.09 (0.30) |
| | slope (stim) | 0.02 (0.15) |
| | correlation | 0.70 |

**Gwei***stim*drug + (stim|cohort:dose:id)

| ***Fixed effects*** | ***b (SE)*** | ***t*** | ***p*** | |
|---|---|---|---|---|
| stim | 0.64 (0.05) | 14.00 | <.01 | * |
| drug | 0.27 (0.10) | 2.66 | 0.01 | * |
| Gwei | 0.36 (0.23) | 1.58 | 0.11 | |
| stim*drug | 0.15 (0.05) | 2.79 | 0.01 | * |
| Gwei *stim | -0.14 (0.15) | -0.97 | 0.34 | |
| Gwei *drug | 0.05 (0.31) | 0.16 | 0.88 | |
| 3-way inter | -0.16 (0.21) | -0.78 | 0.44 | |

| ***Random effects*** | | ***Variance (SD)*** |
|---|---|---|
| cohort:dose:id | intercept | 0.09 (0.30) |
| | slope (stim) | 0.02 (0.15) |
| | correlation | 0.70 |

**n1***stim*drug + (stim|cohort:dose:id)

| ***Fixed effects*** | ***b (SE)*** | ***t*** | ***p*** | |
|---|---|---|---|---|
| stim | 0.64 (0.05) | 13.58 | <0.01 | * |
| drug | 0.23 (0.10) | 2.35 | 0.02 | * |
| n1 | 0.07 (0.01) | 4.63 | <0.01 | * |
| stim*drug | 0.16 (0.06) | 2.97 | 0.01 | * |
| n1 *stim | 0.03 (0.01) | 1.99 | 0.04 | * |

| | | | |
|---|---|---|---|
| n1 *drug | -0.01 (0.02) | -0.80 | 0.42 |
| 3-way inter | -0.01 (0.02) | -0.52 | 0.60 |

| ***Random effects*** | | ***Variance (SD)*** |
|---|---|---|
| cohort:dose:id | intercept | 0.09 (0.29) |
| | slope (stim) | 0.02 (0.16) |
| | correlation | 0.56 |

**n2(U)***stim*drug + (stim|cohort:dose:id)

| ***Fixed effects*** | ***b (SE)*** | ***t*** | ***p*** | |
|---|---|---|---|---|
| stim | 0.64 (0.05) | 13.56 | <0.01 | * |
| drug | 0.24 (0.10) | 2.37 | 0.02 | * |
| n2(U) | 0.10 (0.02) | 4.25 | <0.01 | * |
| stim*drug | 0.17 (0.06) | 3.02 | 0.01 | * |
| n2(U) *stim | 0.04 (0.02) | 0.40 | 0.69 | |
| n2(U) *drug | -0.03 (0.03) | -1.04 | 0.30 | |
| 3-way inter | 0.01 (0.03) | 0.10 | 0.92 | |

| ***Random effects*** | | ***Variance (SD)*** |
|---|---|---|
| cohort:dose:id | intercept | 0.09 (0.29) |
| | slope (stim) | 0.02 (0.16) |
| | correlation | 0.57 |

**n2(W)***stim*drug + (stim|cohort:dose:id)

| ***Fixed effects*** | ***b (SE)*** | ***t*** | ***p*** | |
|---|---|---|---|---|
| stim | .64 (.05) | 13.76 | <0.01 | * |
| drug | .24 (.10) | 2.38 | 0.02 | * |
| n2(W) | .01 (.03) | 0.19 | 0.85 | |
| stim*drug | .17 (.05) | 3.12 | <0.01 | * |
| n2(W) *stim | -.01 (.02) | -0.60 | 0.55 | |
| n2(W) *drug | .04 (.03) | 1.26 | 0.21 | |
| 3-way inter | .07 (.03) | 2.30 | 0.02 | * |

| ***Random effects*** | | ***Variance (SD)*** |
|---|---|---|
| cohort:dose:id | intercept | .09 (.29) |
| | slope (stim) | .02 (.15) |
| | correlation | 0.56 |

**n2wei***stim*drug + (stim|cohort:dose:id)

| ***Fixed effects*** | ***b (SE)*** | ***t*** | ***p*** | |
|---|---|---|---|---|
| stim | .63 (.05) | 13.67 | <0.01 | * |
| drug | .24 (.10) | 2.43 | 0.02 | * |
| n2wei | -.15 (.07) | -2.19 | 0.03 | * |
| stim*drug | .17 (.05) | 3.12 | <0.01 | * |
| n2wei *stim | -.06 (.07) | -0.94 | 0.35 | |
| n2wei *drug | .19 (.08) | 2.38 | 0.02 | * |
| 3-way inter | .19 (.08) | 2.50 | 0.01 | * |

| ***Random effects*** | | ***Variance (SD)*** |
|---|---|---|
| cohort:dose:id | intercept | .09 (.30) |
| | slope (stim) | .02 (.15) |
| | correlation | 0.59 |

**n3(U)***stim*drug3 + (stim|cohort:dose:id)

| ***Fixed effects*** | ***b (SE)*** | ***t*** | ***p*** |
|---|---|---|---|

**n3(W)***stim*drug3 + (stim|cohort:dose:id)

| ***Fixed effects*** | ***b (SE)*** | ***t*** | ***p*** |
|---|---|---|---|

**n3wei***stim*drug3 + (stim|cohort:dose:id)

| ***Fixed effects*** | ***b (SE)*** | ***t*** | ***p*** |
|---|---|---|---|

| | | | | | | | | | | | | | | |
|---|---|---|---|---|---|---|---|---|---|---|---|---|---|---|
| stim | .64 (.05) | 13.61 | <0.01 | * | stim | .64 (.05) | 14.05 | <0.01 | * | stim | .63 (.05) | 13.58 | <0.01 | * |
| drug | .23 (.10) | 2.33 | 0.02 | * | drug | .24 (.10) | 2.42 | 0.02 | * | drug | .26 (.10) | 2.53 | 0.01 | * |
| n3(U) | .11 (.03) | 4.04 | <0.01 | * | n3(W) | -.05 (.04) | -1.40 | 0.16 | | n3wei | -.36 (.09) | -3.89 | <0.01 | * |
| stim*drug | .16 (.06) | 2.97 | 0.01 | * | stim*drug | .16 (.05) | 3.10 | <0.01 | * | stim*drug | .17 (.05) | 3.12 | <0.01 | * |
| n3(U) *stim | .07 (.03) | 2.70 | 0.01 | * | n3(W) *stim | .03 (.03) | 0.90 | 0.37 | | n3wei *stim | -.04 (.09) | -0.44 | 0.66 | |
| n3(U) *drug | -.04 (.03) | -1.27 | 0.21 | | n3(W) *drug | .09 (.04) | 2.10 | 0.04 | * | n3wei *drug | .34 (.11) | 3.07 | <0.01 | * |
| 3-way inter | -.04 (.03) | -1.21 | 0.23 | | 3-way inter | .02 (.04) | 0.59 | 0.56 | | 3-way inter | .15 (.10) | 1.46 | 0.14 | |
| ***Random effects*** | | | ***Variance (SD)*** | | ***Random effects*** | | | ***Variance (SD)*** | | ***Random effects*** | | | ***Variance (SD)*** | |
| cohort:dose:id | intercept | | 0.09 (0.29) | | cohort:dose:id | intercept | | .09 (.29) | | cohort:dose:id | intercept | | .09 (.30) | |
| | slope (stim) | | 0.02 (0.16) | | | slope (stim) | | .02 (.15) | | | slope (stim) | | .02 (.15) | |
| | correlation | | 0.57 | | | correlation | | 0.56 | | | correlation | | 0.59 | |

*Notes*. G = Global prior, n1 / n2 / n3 = Local priors, (U) = Unweighted prior, (W) = Precision-weighted prior, '…wei' = Prior precision weights, stim = stimulus, t = t-values, p = p-values, SE = Standard error, SD=Standard deviation, corr=correlation. * $p$<0.05, ᴼ 0.05<$p$<0.09 (marginally significa
